# scPerturb: Harmonized Single-Cell Perturbation Data

**DOI:** 10.1101/2022.08.20.504663

**Authors:** Stefan Peidli, Tessa D. Green, Ciyue Shen, Torsten Gross, Joseph Min, Samuele Garda, Bo Yuan, Linus J. Schumacher, Jake P. Taylor-King, Debora S. Marks, Augustin Luna, Nils Blüthgen, Chris Sander

## Abstract

Recent biotechnological advances led to growing numbers of single-cell perturbation studies, which reveal molecular and phenotypic responses to large numbers of perturbations. However, analysis across diverse datasets is typically hampered by differences in format, naming conventions, and data filtering. In order to facilitate development and benchmarking of computational methods in systems biology, we collect a set of 44 publicly available single-cell perturbation-response datasets with molecular readouts, including transcriptomics, proteomics and epigenomics. We apply uniform pre-processing and quality control pipelines and harmonize feature annotations. The resulting information resource enables efficient development and testing of computational analysis methods, and facilitates direct comparison and integration across datasets. In addition, we introduce E-statistics for perturbation effect quantification and significance testing, and demonstrate E-distance as a general distance measure for single cell data. Using these datasets, we illustrate the application of E-statistics for quantifying perturbation similarity and efficacy. The data and a package for computing E-statistics is publicly available at scperturb.org. This work provides an information resource and guide for researchers working with single-cell perturbation data, highlights conceptual considerations for new experiments, and makes concrete recommendations for optimal cell counts and read depth.

## Introduction

### [Definition of single-cell perturbation data]

Perturbation experiments probe the response of cells or cellular systems to changes in conditions. These changes traditionally acted equally on all cells in the model system, such as by modifying temperature or adding drugs. Nowadays, with the latest functional genomics techniques, single-cell genetic perturbations which act on individual cellular components have become available. Perturbations using different technologies target different layers of the hierarchy of protein production (Fig 1). At the lowest layer, CRISPR-cas9 acts directly on the genome, using indels to induce frameshift mutations which effectively knock out one or multiple specified genes (Datlinger et al., 2017; Dixit et al., 2016; Jaitin et al., 2016). Newer CRISPRi and CRISPRa technologies inhibit or activate transcription respectively (Gilbert et al., 2014). CRISPR-cas13 acts on the next layer in the hierarchy of protein production to promote RNA degradation (Wessels et al., 2022). Most small molecule drugs, in contrast, act directly on protein products like enzymes and receptors and can have inhibitory or activating effects. When these techniques are applied to large-scale screens, they create a map between genotype, transcriptome, protein, chromatin accessibility, and in some cases phenotype (Frangieh et al., 2021). Single cells are perturbed using unique CRISPR guides, and their corresponding individual barcodes are read out alongside scRNA-seq, CITE-seq or scATAC-seq reads to identify each cell’s perturbation condition (Adamson et al., 2016; Dixit et al., 2016; Frangieh et al., 2021; Rubin et al., 2019). Sequencing with multi-omic readout has been applied to perturbation experiments only recently. CITE-seq, which assesses surface protein counts using oligonucleotide-tagged antibodies measured alongside the transcriptome, has been applied successfully as Perturb-CITE-seq (Frangieh et al., 2021). Efforts have also been made to link Perturb-seq to an ATAC-seq readout (Rubin et al., 2019).

**Fig 1:**
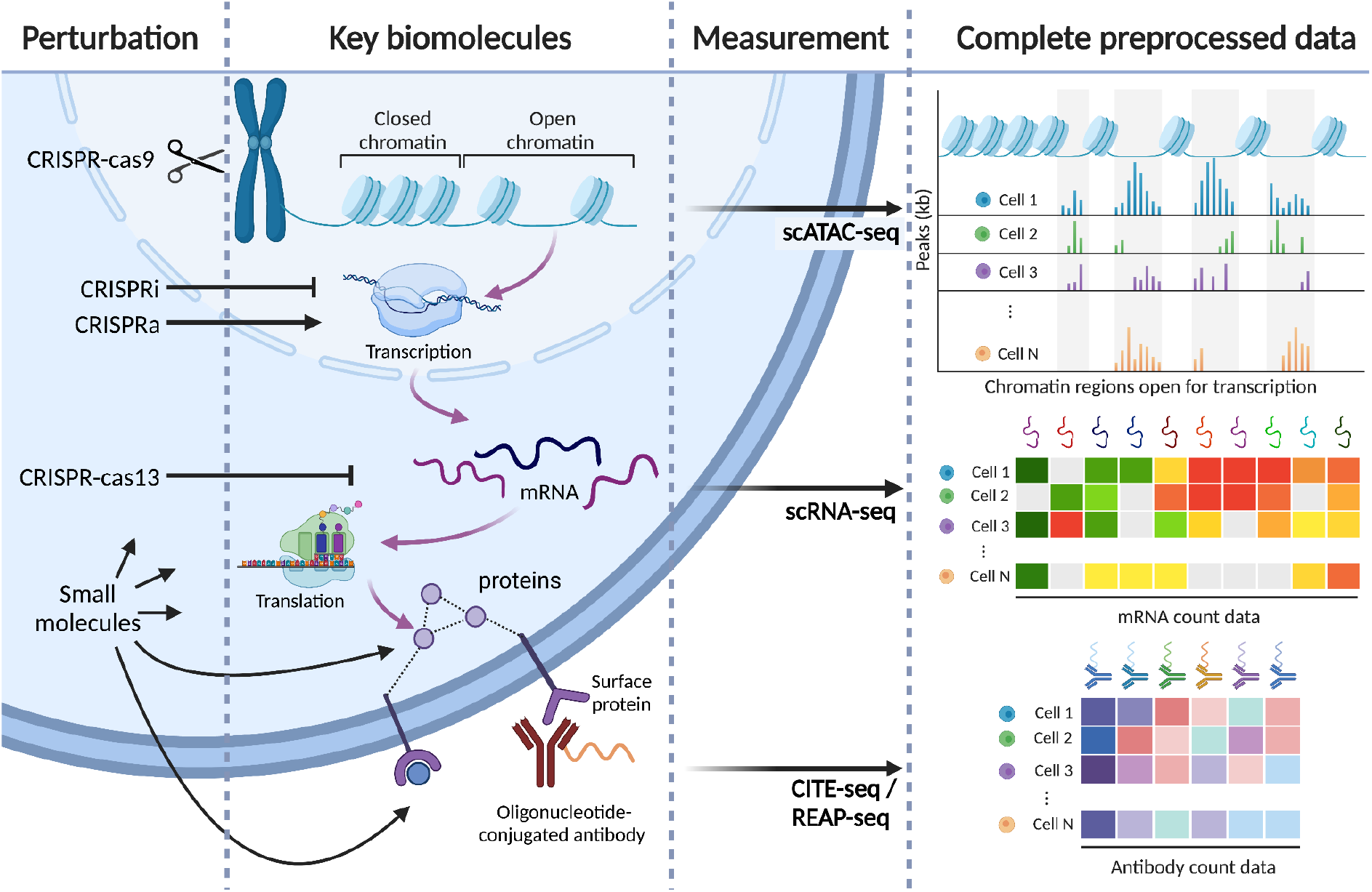
Perturbation-response profiling for single cells. Different perturbations act at different layers in the hierarchy of gene expression and protein production (purple arrows). Perturbations included in scPerturb include CRISPR-cas9, which directly perturbs the genome; CRISPRa, which activates transcription of a target gene; CRISPRi, which blocks transcription of targeted genes; CRISPR-cas13, which cleaves targeted mRNAs and promotes their degradation; cytokines that bind cell surface receptors; and small molecules perturb various cellular mechanisms. Single cell measurements probe the response to perturbation, also at different layers of gene expression: scATAC-seq directly probes chromatin state; scRNA-seq measures mRNA; and protein count data currently is typically obtained via antibodies bound to proteins.

### [Uses of single-cell perturbation data]

Large-scale single-cell perturbation-response screens enable exploration of complex cellular behavior not accessible from bulk observation. Directionality in regulatory network models cannot be inferred without interventional or time-series data about the system (Gross et al., 2019). Experiments with targeted perturbations can be modeled as affecting individual nodes of a regulatory network model, while the molecular readouts provide information on the state changes. This creates the opportunity to investigate mechanistic processes and infer regulatory interactions and their directionality (Pratapa et al., 2020). Typically, however, perturbation datasets have still been too small to elucidate the complexity of a cellular system, and thus accurately predictive models of regulatory interactions remain difficult to infer (Gross and Blüthgen, 2020). This limitation will be reduced as dataset size continues to increase. More directly, drug screens have been used to suggest therapeutic interventions by analyzing detailed molecular effects of targeted drugs, and designing new single or combinations of perturbations (Bertin et al., 2022; Franz et al., 2021; Preuer et al., 2018).

### [Motivation for a distance measure for high-dimensional expression profiles]

Reliable analysis of increasingly large perturbation datasets requires statistical tools powerful and efficient enough to harness both the massive number of cells and perturbations and the inherently high dimensionality of the data. This high dimensionality complicates calculation of distances between perturbations, as does cell-cell variation and data sparsity (Kharchenko, 2021). There is not presently a convention for the statistical comparison measure used in perturbation studies. Some studies calculate pseudo-bulk by combining all cells in a given perturbation (Adamson et al., 2016; Datlinger et al., 2017). This means losing any information about the variation within each cell type. Studies with mixtures of cell types do the opposite, developing complex methods for quantifying similarity between heterogeneous cell populations (Burkhardt et al., 2021; Dann et al., 2022; Gehring et al., 2020). Ideally, one would perform statistical comparisons to identify similar perturbations and classify perturbation strength using a multivariate distance measure between sets of cells. Such a distance measure describes the difference or similarity between sets of cells treated with distinct perturbations, thus inferring difference or similarity in terms of mechanism or perturbation target; shared mechanisms tend to produce similar shifts in molecular profiles (Replogle et al., 2022; Tian et al., 2021). A number of distance measures for scRNA-seq have been explored by the single-cell community in recent years, including Wasserstein distances (Chen et al., 2020), maximum mean discrepancy (Lotfollahi et al., 2020), neighborhood-based measures (Burkhardt et al., 2021; Dann et al., 2022), E-distance (Replogle et al., 2022), Here we exclusively use the E-distance, a fundamental statistical measure of distances between point clouds that can be used in a statistical test to identify strong or weak perturbations as well as to distinguish between perturbations affecting distinct cellular sub-processes (see Methods). This test is a statistically reliable tool for computational diagnostics of information content for a specific perturbation and can inform design of experiments and data selection for training models.

### [Motivation for unifying datasets]

Each large perturbation screen is specifically designed to study a particular system under a set of perturbations of interest. This results in a heterogeneous assortment of single-cell perturbation-response data with a wide range of different cell types, such as immortalized cell lines and iPSC-derived models, and different perturbation technologies, including knockouts, activation, interference, base editing, and prime editing (Przybyla and Gilbert, 2022). Novel computational methods to efficiently harmonize these different perturbation datasets on a large scale are needed. Such integrative analysis is complicated by batch effects and biological differences between primary tissue and cell culture (Forcato et al., 2021; Luecken et al., 2022). Published computational methods for perturbation data are primarily focused on individual datasets (Duan et al., 2019; Jin et al., 2022; Lotfollahi et al., 2019). Moving from single-dataset to multi-dataset analysis will require development of principled quantitative approaches to perturbation biology; the dataset resource based on this work can serve as a foundation for building these models going forward.

### [Prior work]

While large databases of perturbations with bulk readouts exist, single-cell perturbation technologies are newer and data is still not unified (Stathias et al., 2020; Tsherniak et al., 2017). Existing collections of datasets are primarily a means for filtering datasets but do not supply a unified format for perturbations. Some of these collections of single-cell datasets were produced to benchmark computational methods for data integration (Lance et al., 2022). Another collection specifically aggregated all available data in the single-cell literature, with well over a thousand datasets described in a sortable table, but with no attempt at data unification and labels focused on observational studies (Svensson et al., 2020). The Broad Institute’s Single Cell Portal provides .h5 files for 478 studies with cell type names from a common list but does not harmonize the datasets or allow for filtering by perturbation (Broad Institute, 2022). Yet, unified datasets are key for developing generalizable machine learning methods and establishing multimodal data integration. A recent review and repository of single-cell perturbation data for machine learning lists 22 datasets but supplied cleaned and format-unified data for only 6 (Ji et al., 2021). An existing unified framework for single cell data, called ‘sfaira’, is ideal for model building and memory efficient data loading, but the public ‘data zoo’ does not currently supply perturbation datasets or standardized perturbation annotations (Fischer et al., 2021).

### [Our contribution]

We aim to provide a resource of standardized datasets reporting targeted perturbations with single-cell readouts and to facilitate the development and benchmarking of computational approaches in system biology. We collected a set of 44 publicly available perturbation-response datasets from 25 papers (Table 1, Supp Fig 3B). Our perturbation strength quantification and comparison of perturbation-specific variables, such as the number of perturbations and the number of cells per perturbation, across experiments may serve as a reference for optimal experimental design of future single-cell perturbation experiments. We also describe the E-distance and E-test as tools for statistical comparisons of sets of cells and benchmark their robustness and applicability for distinguishing both perturbations and cell types across different datasets and modalities. A web interface for data access, analysis and visualization is available at scperturb.org, and a Python implementation of e-distance based statistics for single cell data is publicly available as `scperturb` on PyPI.

**Table 1:** Key metadata for datasets on scPerturb.org. More details in Supp Table 1.

| Source Paper | Modality | Perturbation type | Number of perturbations |
| --- | --- | --- | --- |
| (Adamson et al., 2016) | RNA | CRISPRi | 9, 20, 114 |
| (Aissa et al., 2021) | RNA | drugs | 4 |
| (Chang et al., 2022) | RNA | drugs | 4 |
| (Datlinger et al., 2017) | RNA | CRISPR-cas9+TCR<sup>&</sup> | 97 |
| (Datlinger et al., 2021) | RNA | CRISPR-cas9+TCR<sup>&</sup> | 48 |
| (Dixit et al., 2016) | RNA | CRISPR-cas9 | 31 |
| (Frangieh et al., 2021) | RNA + protein | CRISPR-cas9 | 249 |
| (Gasperini et al., 2019) | RNA | CRISPRi | 43314*, 39087*, 16531* |
| (Gehring et al., 2020) | RNA | drugs | 4 |
| (Liscovitch-Brauer et al., 2021) | ATAC | CRISPR-cas9 | 22,84 |
| (McFarland et al., 2020) | RNA | drugs, CRISPR-cas9 | 18 |
| (Mimitou et al., 2021) | ATAC + protein | CRISPR-cas9 | 6 |
| (Norman et al., 2019) | RNA | CRISPRa | 237 |
| (Papalexi et al., 2021) | RNA + protein | CRISPR-cas9 | 11,99 |
| (Pierce et al., 2021) | ATAC | CRISPRi | 41,41,41 |
| (Replogle et al., 2022) | RNA | CRISPRi | 2058, 2394, 9867 |
| (Schiebinger et al., 2019) | RNA | cytokines | 2,3 |
| (Schraivogel et al., 2020) | RNA | CRISPR-cas9 | 3105*, 4115* |
| (Shifrut et al., 2018) | RNA | CRISPR-cas9+TCR<sup>&</sup> | 49 |
| (Srivatsan et al., 2020) | RNA | drugs | 5, 8, 189 |
| (Tian et al., 2019) | RNA | CRISPRi | 27 |
| (Tian et al., 2021) | RNA | CRISPRa, CRISPRi | 101, 185 |
| (Weinreb et al., 2020) | RNA | cytokines | 5 |
| (Xie et al., 2017) | RNA | CRISPR-cas9 | 229 |
| (Zhao et al., 2021) | RNA | drugs | 7 |

\*: perturbation total treats perturbations A, B, and (A and B) as three unique perturbations
&: TCR receptor stimulation

## Results

### [Overview of datasets in the information resource]

Molecular readouts for our 44 publicly available single-cell perturbation response datasets include transcriptomes, proteins and epigenomes (Table 1, Fig 2A). Metadata was harmonized across datasets (Supp Table 2). 32 datasets in this resource were perturbed using CRISPR and 9 datasets perturbed with drugs. This paucity of drug datasets is likely due to the experimental hurdle of applying large numbers of perturbations to cells; although it is possible to set up multiplexed sequencing for arrayed treatment conditions, this entails a large amount of manual labor necessary to set up hundreds of separate wells with individual drugs, limiting the total number of drug perturbations in a single experiment. In contrast, the mixed set of single guides for CRISPR perturbations can be applied in parallel, allowing these experiments to be scaled up massively. While 32 datasets measure scRNA-seq exclusively, we also include scATAC from three papers, including one with simultaneous protein measurements (Mimitou et al., 2019). For each scRNA-seq dataset we supply count matrices, where each cell has a perturbation annotation, quality control metrics including gene counts and mitochondrial read percentage. Quality control plots for each dataset are also available on scperturb.org (for an example see Supp Fig 1). Three CITE-seq datasets are included with protein and RNA counts separately downloadable (Frangieh et al., 2021; Papalexi et al., 2021).

**Fig 2:**
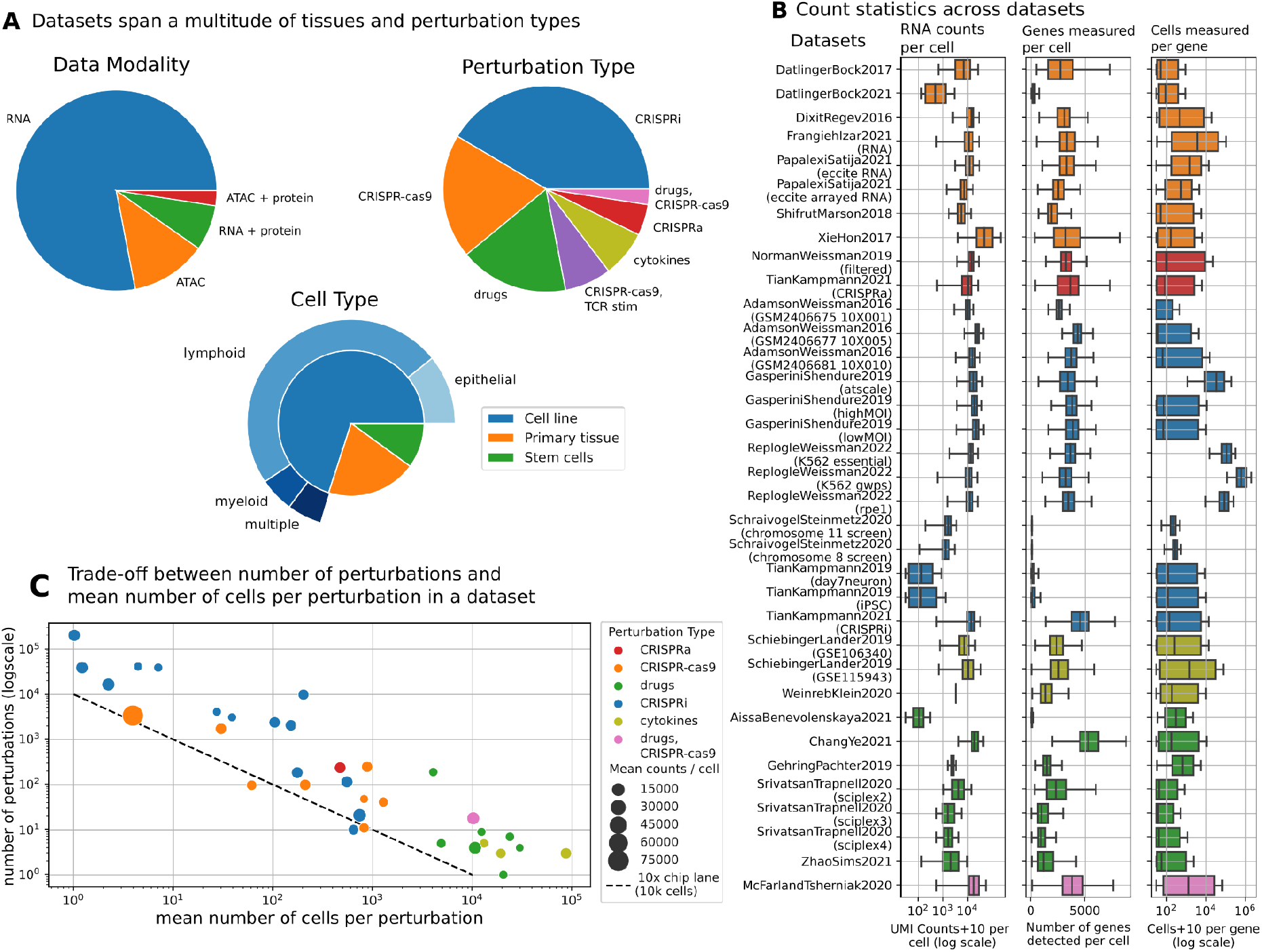
Single cell perturbation-response datasets are diverse in type, size, and quality. (A) The majority of included datasets result from CRISPR (DNA cut, inhibition or activation) perturbations using cell lines derived from various cancers. The studies performed on cells from primary tissues generally use drug perturbations. Primary tissue refers to samples taken directly from patients or mice, sometimes with multiple cell types. (B) Sequencing and cell count metrics across scPerturb perturbation datasets (rows), colored by perturbation type as in Figure 2C. From left to right: Distribution of total RNA counts per cell (left); distribution of the number of genes with at least one count in a cell (middle); distribution of number of cells with at least one count of a gene per gene (right). Most datasets have on average approximately 3000 genes measured per cell, though some outlier datasets have significantly sparser coverage of genes. (C) Each circle represents one dataset. Due to experimental constraints, most datasets have approximately the same number of total cells, pooled across a set of marked perturbations, resulting in a tradeoff between the number of perturbations and the number of cells in each perturbation. CRISPR-perturbation datasets, compared to drug-perturbation datasets, have fewer cells per perturbation but a larger number of perturbations. Due to experimental constraints, most datasets have approximately the same number of total cells, pooled across a set of marked perturbations, resulting in a tradeoff between the number of perturbations and the number of cells in each perturbation.

### [Choice of features in scATAC-seq data]

In contrast to scRNA-seq data, which can be represented naturally as counts per gene, there is no single obvious feature that could be computed for scATAC-seq data, which, in its raw form, provides a noisy and very sparse description of chromatin accessibility over the entire genome. We therefore generated five different feature sets for scATAC-seq data, as motivated in prior studies (Chen et al., 2019; Granja et al., 2021). These either attempt to summarize chromatin accessibility information over different types of biologically relevant genomic intervals (e.g. gene neighborhood), or represent dense low-dimensional embeddings of the original data (see Methods for details). Different features address different biological questions in different contexts (Pierce et al., 2021), which is why we include all five feature sets in our resource.

### [Count statistics in scRNA-seq]

Sample quality measures vary significantly across datasets (Fig 2B). Total unique molecular identifier (UMI) counts per cell and number of genes per cell are calculated as described in (Luecken and Theis, 2019). The average sequencing depth, i.e. the mean number of reads per cell, in each study affects the number of lowly expressed genes observed. Increasing the sequencing depth increases the number of UMI counts measured even for lowly expressed genes, reducing the uncertainty associated with zero counts (Haque et al., 2017; Svensson et al., 2020). These differences can affect the distinguishability of perturbations and performance of downstream analysis methods.

### [Cells per perturbation]

The total number of cells per dataset is usually restricted by experimental limitations, though has increased over time (Supp Fig 2A). Therefore, there is a tradeoff between the number of perturbations and the mean number of cells per perturbation in a dataset (Fig 2C). The type of perturbation partially dictates the number of cells per perturbation; CRISPR datasets tend to have more perturbations than drug datasets because they are easier to scale up using multiplexing, with a corresponding smaller number of cells per perturbation. This is visible in Figure 2B, where CRISPR datasets are clustered to the left of the plot.

### [E-distance: definition]

To compare and evaluate perturbations within each dataset we utilized the E-distance, a statistical distance measure between two distributions, which was used as a test statistic in (Replogle et al., 2022). Essentially, the E-distance compares the mean pairwise distance of cells across two different perturbations to the mean pairwise distance of cells within the two distributions (see Methods). If the former is much larger than the latter, the two distributions can be seen as distinct. Similar to Replogle et al., we compute the E-distance after PCA (Principal Component Analysis) for dimensionality reduction (see Methods).

### [E-distance: interpretation]

The E-distance provides intuition about the signal-to-noise ratio in a dataset. For two groups of cells, it relates the distance between cells across the groups (“signal”), to the width of each distribution (“noise”) (Fig 3A). If this distance is large, distributions are distinguishable, and the corresponding perturbation has a strong effect. A low E-distance indicates that a perturbation did not induce a large shift in expression profiles, reflecting either technical problems in the experiment, ineffectiveness of the perturbation, or perturbation resistance.

**Fig 3:**
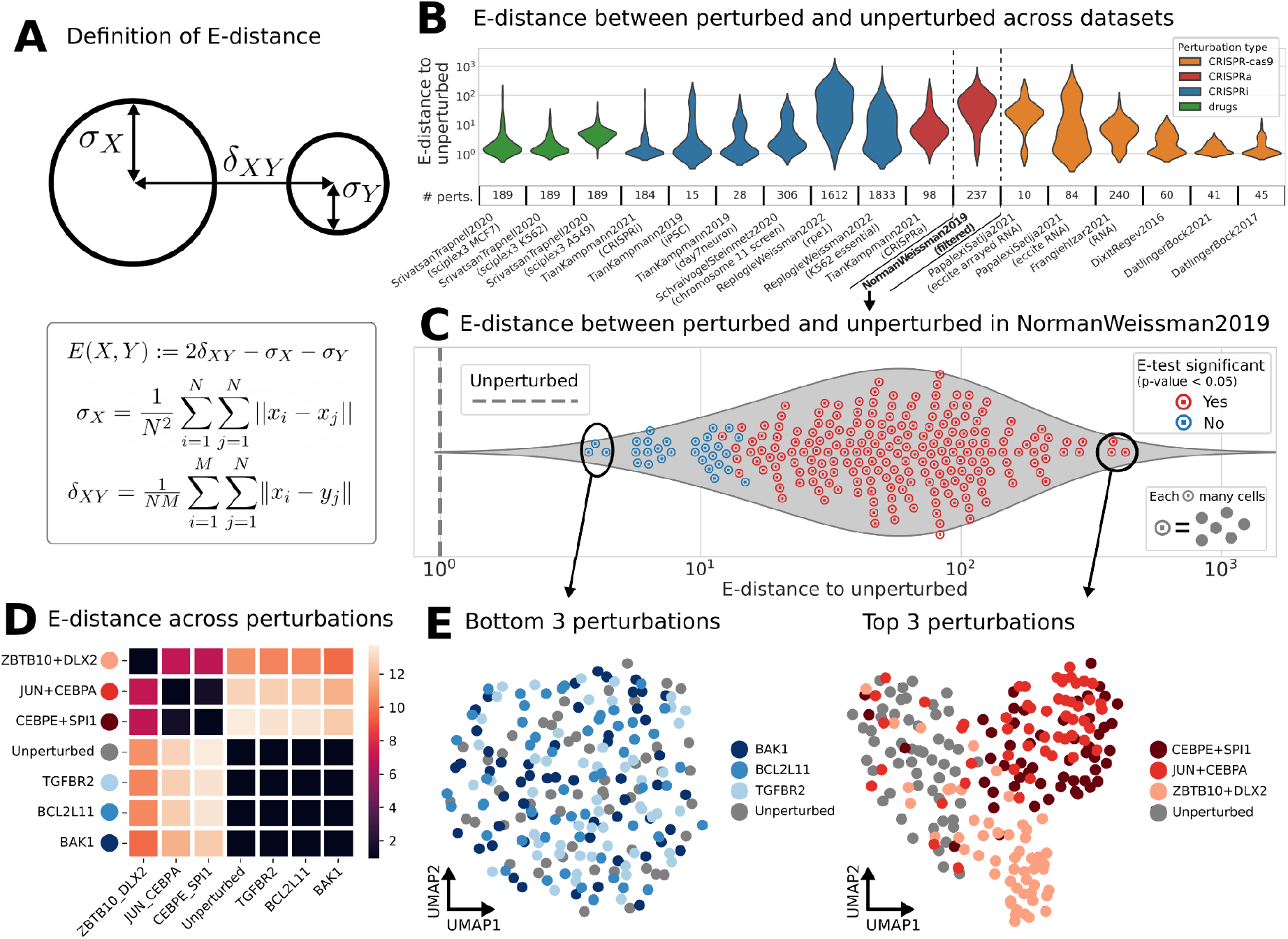
E-statistics describe distinctiveness of perturbations in single-cell data. (A) Definition of E-distance, relating the width of cell distributions of high-dimensional molecular profiles to their distance from each other (see Methods). A large E-distance of perturbed cells from unperturbed indicates a strong change in molecular profile induced by the perturbation. (B) Distribution of E-distances (plus 1 for log scale, same in Fig 3C) between perturbed and unperturbed cells across datasets. The number of perturbations per dataset is displayed along the bottom. Note that this plot is best used to compare the shape of the E-distance distribution rather than the magnitude; the mean E-distance will vary significantly with other dataset properties. (C-E) Analysis based on E-statistics for one selected dataset (Norman et al., 2019): (C) Distribution of E-distances between perturbed and unperturbed cells as in Figure 3B. Each circled point is a perturbation, i.e., represents a set of cell profiles. Each perturbation was tested for significant E-distance to unperturbed (E-test). (D) Pairwise E-distance matrix across the top and bottom 3 perturbations of Figure 3C and the unperturbed cells. (E) UMAP of single cells of the weakest (left, Bottom 3) and strongest (right, Top 3) perturbations.

### [E-distance: distinguishing cell types]

To test whether E-distance values replicate differences between well-known cell types, we applied the E-distance to a CITE-seq human PBMC (peripheral blood mononuclear cells) dataset with existing cell type annotations (Hao et al., 2021). Separately for RNA and protein (from antibody-derived tags), we computed PCA-based E-distances between all pairs of cell types, equivalent to how perturbation E-distance is computed. The resulting pairwise distance matrices were used to compute cell type hierarchies (see Methods), which we compare to known cell type relationships (Fig 4A, 4B, Diehl et al., 2016). In both data modalities, B cell subtypes are clustered together, and platelets are the most distinct from any other cell type. Lymphoid and myeloid cells form two separate groups in the E-distance hierarchy. Notably, innate lymphoid cells (ILCs) and NK cell clusters form a distinct group as well. ILCs are innate immune cells that functionally correspond to specific types of classical lymphocytes properly expressing diversified antigen receptors; NK cells are a type of ILC also known as ILC4 and are functionally similar to cytotoxic T cells (Artis and Spits, 2015; Vivier et al., 2018). This functional similarity translates to strong similarities in transcriptional profiles, which often leads to difficulties in distinguishing NK cells and cytotoxic T cells in scRNA-seq data. These cell types are more easily disentangled by protein marker based distances, exemplifying the usefulness of CITE-seq as a method for identifying immune cell types. Likewise, when using protein, T cells are clustered primarily by CD4/CD8 type, whereas, when using RNA, they are clustered by functional phenotype (naive, proliferating, memory). For instance, clustering the cells with the E-distance in RNA-space separates proliferating cells of many types into a single cluster, likely due to shared expression of cell-cycle related genes; the cell cycle is known to have a strong effect on the transcriptome profile of cells and is not captured by surface protein measurements. We conclude that, by comparing RNA and protein representations, the protein modality more accurately represents cell type differences traditionally defined by immunologists on the basis of surface proteins, whereas the RNA representation primarily reflects functional programs of the cells such as cytotoxicity or proliferation. In both cases, the E-distance accurately captures known characteristics of each measurement modality.

**Fig 4:**
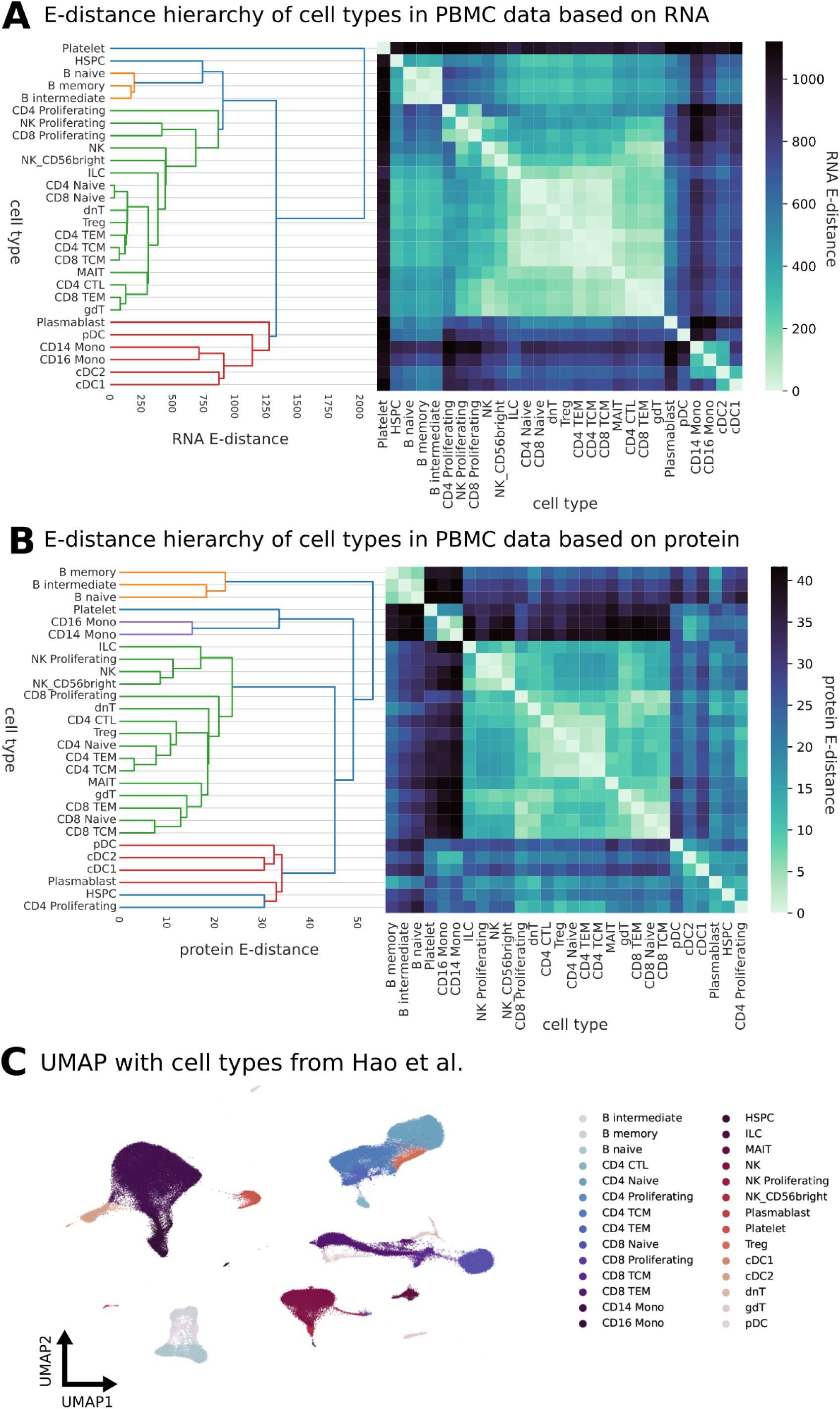
Cell type hierarchies computed using E-distance match known cell type relationships. (A) Hierarchical clustering of pairwise E-distances computed using RNA matches prior knowledge of transcriptome-defined cell types. Dendrogram and heatmap use the same distances. Data from (Hao et al., 2021). (B) As in (A) but using antibody-tagged surface proteins instead of RNA. (C) Visualization of cell type relationships in full multimodal dataset after batch correction. Coordinates and cell type annotations from (Hao et al., 2021).

### [E-distance: E-test]

The E-distance can also be used as a test statistic to assess whether cells after a perturbation are significantly different from unperturbed cells (Replogle et al., 2022), Supp Table 3). The E-test is a permutation test that uses the E-distance as a test statistic (Székely and Rizzo, 2013, details in Methods). This permutation test requires hundreds of iterations of computing the E-distance on randomized data, but is necessary for direct comparisons of perturbations across studies, as the exact value of the E-distance depends on dataset-specific parameters. The lack of robustness is exemplified by Figure 5A, which indicates that reducing the number of cells actually increases the E-distance, while the E-test gradually loses significance. Thus, while the E-distance is a useful tool for analysis within one dataset or between experimentally similar datasets, we recommend the E-test as the appropriate statistical measure for comparisons of perturbation effects between different datasets.

**Fig 5:**
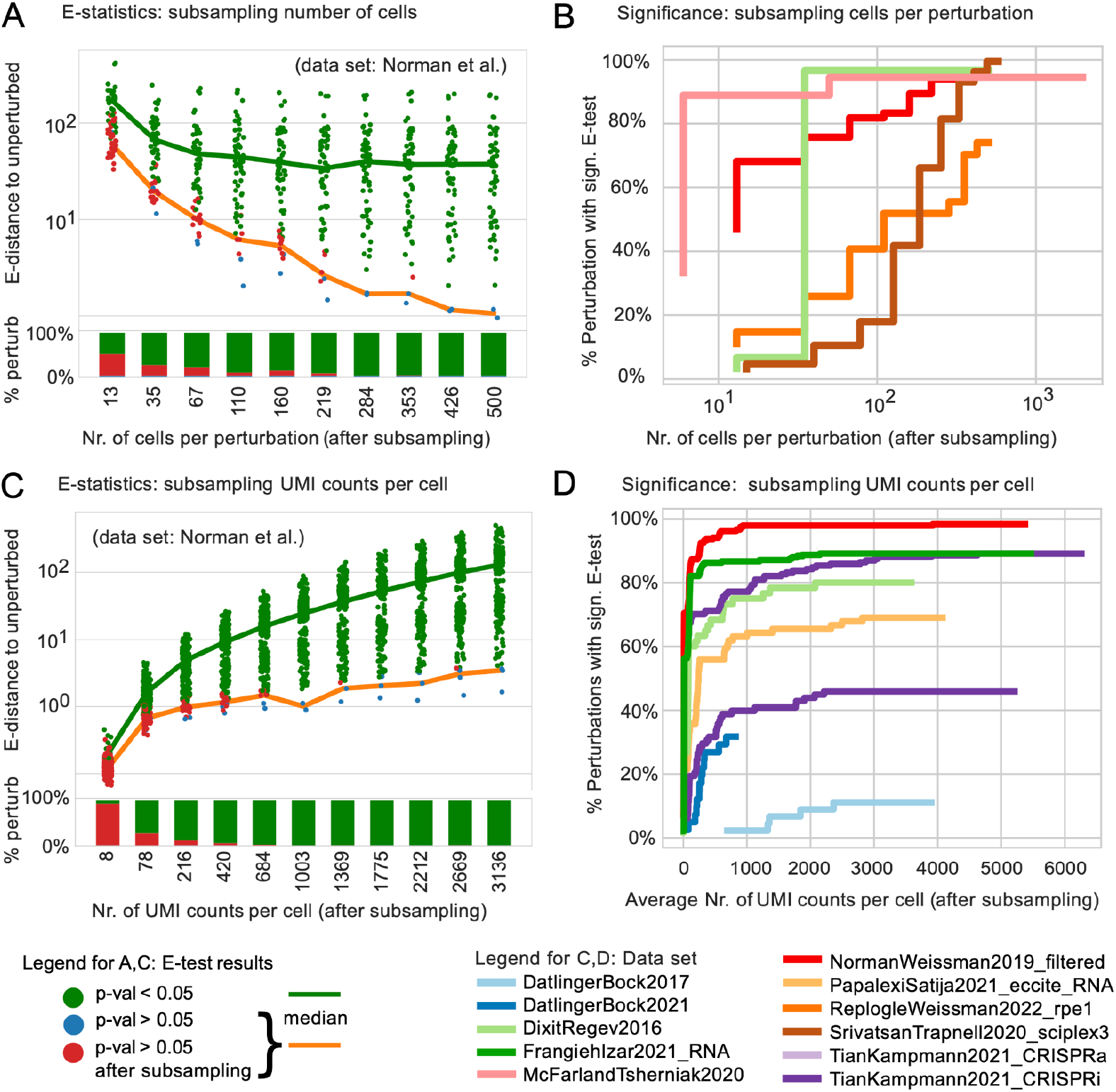
Effect of subsampling UMI counts per cell and number of cells per perturbation on E-statistics. (A) E-distance of each perturbation to unperturbed in Norman et al. while subsampling the number of cells per perturbation; Color indicates E-test results; “significance lost”: perturbation significant when all cells are considered, but not significant after subsampling. The E-test loses significance with lower cell numbers while the E-distance actually increases. (B) Overall number of perturbations with significant E-test decreases when subsampling cells. (C) As in Figure 5A but subsampling UMI counts per cell while keeping the number of cells constant. Loss of E-test significance and dropping E-distance to unperturbed as overall signal gets deteriorated with removal of UMI counts. (D) As in Figure 5B but subsampling UMI counts per cell while keeping the number of cells constant.

### [E-distance: perturbation dataset example]

Interestingly, we found that E-distances between perturbed and unperturbed cells vary significantly across datasets (Fig 3B). The dataset labeled with “NormanWeissman2019” had the largest mean E-distance between all perturbations (Norman et al., 2019) compared to datasets of similar size. In fact, expression profiles of most perturbations in this dataset were significantly different from those of unperturbed cells according to the E-test (Fig 3C). Plausibly, this is in part caused by two-target perturbations using CRISPRa in that dataset: targeting the same gene with two single guides increases the chances of causing a considerable change in the transcript profile. Indeed, the three perturbations with highest E-distance are double perturbations while the three closest in E-distance are not. The corresponding UMAPs for these perturbations, computed using the same PCs as the E-distance, provide a confirmatory visual intuition for high and low E-distances (Fig 3E). The top three perturbations causing the largest E-distance to unperturbed are easily distinguishable from the gray unperturbed cells, while the bottom three weakest perturbations are part of a single, uniform cloud virtually indistinguishable from the unperturbed cells. The smallest E-distance thus results from perturbations which have the least effect on the distribution of cells.

### [E-distance: perturbation-perturbation distances]

The E-distance can also be used to measure similarity between different perturbations. For instance, there is a clear overlap of CEBPE+SPI1 and JUN+CEBPA perturbed cells in the UMAP (Fig 3E). This overlap is captured by the low E-distance between the two perturbations; these two perturbations are closer to each other than they are to unperturbed cells or to other perturbations (Fig 3D, Supp Fig 3A). We envision that the E-distance can be used as a suitable distance for other downstream tasks such as drug embeddings and clustering of perturbations, which could allow inference of functional similarity of perturbations by similarity in their induced molecular responses measured by the E-distance.

### [E-distance: Effect of number of cells on the E-statistics]

We investigate the robustness of E-distance and E-test scores to experimental and computational parameters using our extensive collection of harmonized single-cell perturbation datasets. We subsampled the number of cells per perturbation to create artificially smaller datasets, then examined how the E-distance and E-test results change. We find that the E-distance actually increases as the number of cells per perturbation decreases, indicating that cells per perturbation should be standardized prior to calculating E-distances (Fig 5A). This increase reflects the fact that the E-distance, as a V-statistic, is a biased estimator (Székely and Rizzo, 2013). Despite the increase in E-distance with falling cell numbers, the number of significant perturbations correctly decreases with fewer cell counts, and only some datasets have saturated significance at full number of cells in that dataset (Fig 5B). This saturation point will depend on the strength of the perturbation and on the heterogeneity of the dataset; if all cells are similar to each other, a small set of cells will sufficiently describe every possible response to a perturbation. This suggests that, unsurprisingly, increasing sample size enables discovery of significant perturbations with smaller magnitude.

### [E-distance: Effect of number of UMI counts on the E-statistics]

Similarly, we subset the number of UMI (unique molecular identifier) counts per cell, finding that E-distance increases as the number of UMI counts per cell increases (Fig 5C). The number of significant perturbations under the E-test, though, saturates around 500 counts per cell, with most perturbations that were significant at the full measured read depth maintaining that significance even with far fewer counts per cell (Fig 5D). The stability of E-test results with respect to UMI counts, in contrast to the actual E-distance value, exemplifies the necessity of the E-test as the appropriate statistical measure to evaluate perturbation effects. The optimal UMI and cell counts for a given experiment depend on downstream specific modeling tasks, as discussed in more detail elsewhere (Gross and Blüthgen, 2020). As a baseline for significant perturbations, as defined by the E-test, we suggest at least 300 cells per perturbation (Fig 5B) and 1000 average UMI counts per cell (Fig 5D) as an experimental guideline for distinguishable perturbations.

### [E-distance: robust to calculation choices]

We also examined how choices made in computing the principal components (PCs) used in distance calculations affects E-statistics. The number of PCs used from PCA to compute the E-distance had a moderate effect on E-test results, mildly decreasing the number of significant perturbations (Supp Fig 5A). Interestingly, E-test significance was lost most rapidly in a TAP-seq (targeted perturb-seq) experiment (Schraivogel et al., 2020). TAP-seq only measures approximately 3000 genes of interest, and thus has far fewer starting features than other datasets. This leads to reduced correlation between genes in the resulting expression matrix, and thus fewer PCs are needed to sufficiently describe the data. The number of highly variable genes (HVGs) used to compute the PCs had almost no effect on E-testing above 500 HVGs (Supp Fig 5B). Computing PCs separately for each perturbation rather than jointly across all perturbations in a given dataset similarly had minimal impact on the resulting E-distances (Supp Fig 5C). Taken together, this analysis indicates that E-statistics can be calculated as part of an existing computational workflow, which already includes calculating PCs.

### [Dataset highlights]

With this assembly of datasets and quality control metrics described below, we were able to nominate notable datasets. The most extensive drug dataset is the sci-Plex 3 dataset with over 188 drugs tested across three cell lines (Srivatsan et al., 2020); 107 of those perturbations were significant according to E-test analysis (Supp Table 3). Five drugs in this dataset also appear in other drug perturbation datasets (Supp Table 4). We hope that future large-scale drug screens will enable more detailed analysis of drug response across different cell types and conditions. Another drug dataset applies combinations of three drug perturbations at varying concentrations across samples (Gehring et al., 2020). We excluded this dataset from the E-distance analysis due to its complex study design, which was not directly comparable to any other included studies. By far the most detailed CRISPR dataset is from a recently published study which perturbed 9867 genes in human cells (ReplogleWeissman2022). Containing >2.5 million cells, this dataset is the largest in our database, with the number of cells each gene is detected in significantly higher than in other datasets (Fig 2B). Notably, 138 CRISPR perturbations are seen in both RNA and ATAC datasets (Supp Table 5). More than 100 genes perturbed with CRISPRa in one dataset are perturbed with CRISPRi perturbations in another dataset of the same cell line, either in one paper (Tian et al., 2021) or across multiple studies (Norman et al., 2019; Replogle et al., 2022). The most frequently perturbed gene, MYC, is perturbed in 9 datasets from 3 papers. Protein, RNA and ATAC readouts for CRISPRi perturbation of MYC are all available for K562 cells (Frangieh et al., 2021; Pierce et al., 2021; Replogle et al., 2022).

## Discussion

### [Concise summary of results]

We present a dataset resource and an intuitive analytic method for quantifying and analyzing single-cell perturbation datasets. Datasets are described in detail, with additional individualized quality control metrics available on scperturb.org. The uniform annotations in this resource will enable data integration and benchmarking as well as exploration of shared perturbations across datasets. The use of the E-distance is motivated and applied to quantitatively compare perturbations within each dataset. We illustrate how to interpret high and low E-distances and use E-distances to identify functionally similar versus distinct perturbations. We also investigate the effect of dataset specific parameters on E-statistics, showing that E-statistics stabilize above 1000 counts per cell and 300 cells per perturbation.

### [Overlap of perturbations across studies]

While this work simplifies access to datasets, joint analysis of single-cell datasets is limited by the complexity of data integration. Across the eight drug datasets examined in this study, only 5 chemical agents occurred in more than one dataset (Supp Table 4). Shared gene targets are found more often across the CRISPR datasets (Supp Table 5). However, multiplicity of infections and other conditions frequently differ as well, and comparisons are further complicated by distinct perturbation methods (Table 1). The considerable overlap of perturbations across studies makes this a useful resource for benchmarking model generalizability. With more datasets anticipated in the future, we will have the unique opportunity to integrate datasets with more overlapping perturbations and nominate machine learning benchmarks for data integration.

### [Towards standardization]

Lack of standardization in data sharing and processing hampered the creation of this resource. Although many processed datasets were available on the NCBI Gene Expression Omnibus (GEO) (Barrett et al., 2013), there is no standard format for sharing CRISPR barcode assignments and other metadata. Starting analysis from sequencing reads may have improved interoperability of datasets in this resource, but guide assignment procedures and demultiplexing algorithms are specific to experimental setup. For scATAC data, data comparison is made more challenging by the lack of a standard method for feature assignment (see Methods). In particular, scATAC feature assignments specific to CRISPR perturbations, where known locus-of-action could be used to improve feature calls (Chen et al., 2019). In all modalities, many datasets only supplied processed data, or raw data was only available after institutional clearance. Adding more datasets to this resource, or the creation of similar resources in the future, would be much easier if there were standard formats for sharing perturbation data, and, more generally, standard formats for sharing single-cell annotations. We think a community-wide discussion on standardization of such data is urgently needed, as has been done for proteomic data (Gatto et al., 2022).

### [Additional considerations for single cell perturbation experimental design]

Experimental design choices such as the optimal number of cells per perturbation and the sequencing depth for each cell depend on the questions the dataset is intended to answer, and on the strength and uniqueness of the gene expression changes caused by the perturbations (Fig 2B). Unfortunately, it is difficult to ascertain to what extent a low E-distance between perturbed and unperturbed cells in the data is caused by technical noise. Increasing the dose or varying the time between perturbation start and harvesting of the cells may be advisable to increase the signal to noise ratio without sequencing more cells. For perturbation distinguishability as defined by the E-test, regardless of experimental parameters, we find that one should have at least 300 cells per perturbation and an average of 1000 UMIs per cell.

### [Conclusion]

We envision that the scPerturb collection of datasets and the suggested E-statistics analytic framework will be a valuable starting point for the analysis of single-cell perturbation data. The unified annotations and perturbation significance testing across datasets should prove especially useful to the machine learning community for training models on this data. We expect new datasets and new experimental perturbation methods in the future will enable the community to develop novel computational approaches which exploit the increasing amount and complexity of single-cell perturbation data, aiming at the development of increasingly accurate and quantitatively predictive models of cell biological processes and the design of targeted interventions for investigational or therapeutic purposes.

## Data Availability

The website scperturb.org stores harmonized datasets with the following:

- scRNA-seq and antibody-based protein datasets: .h5ad files and .mtx files are available, which can be easily read with python or R scripts.
- scATAC-seq: multiple different feature matrix definitions as separate download options.
- Access details for the original publication for each dataset
- Quality control plots for each dataset
- Filtering, e.g., by readout or type of perturbation
- RNA data at https://zenodo.org/record/7041849 and ATAC data at https://zenodo.org/record/7058382

## Code Availability

Open access source code is at https://github.com/sanderlab/scPerturb/. We compiled a corresponding Python package called `scperturb` for performing E-statistics (E-distance and E-testing) in single-cell data, published on PyPI under https://pypi.org/project/scperturb/.

## Methods

### scATAC-seq

#### Data acquisition

We included scATAC-seq data from three different sources: Spear-ATAC (Pierce et al., 2021), CRISPR-sciATAC (Liscovitch-Brauer et al., 2021), and ASAP-seq (Mimitou et al., 2019). All data that was used in our analysis can be programmatically downloaded with scripts that are provided in our code repository (https://github.com/sanderlab/scPerturb).

scATAC-seq is a biomolecular technique to assess chromatin accessibility within single cells (Buenrostro et al., 2015; Cusanovich et al., 2015). The starting point of our data processing pipeline are BED-like tabular fragment files, in which each line represents a unique ATAC-seq fragment captured by the assay. Each fragment is mapped to a genomic interval and a cell barcode. The goal of our pipeline is to extract standardized features from this information. Those are:

- Embeddings derived from Latent-Semantic-Indexing (LSI) (Cusanovich et al., 2015) with 30 dimensions for each cell (a dimensionality reduction method that is well-suited for the sparsity of the data)
- Gene scores that measure the chromatin accessibility around each gene for each cell (the weighted sum of fragment counts around the neighborhood of a gene’s transcription start site where more distant counts contribute less)
- A peak-barcode matrix that quantifies the chromatin accessibility at (data-set specific) consensus peaks (genomic intervals) for each cell
- ChromVar scores (Schep et al., 2017), which quantify the activity of a set of transcription factors for each cell, using transcription factor footprints as defined in (Vierstra et al., 2020)
- Marker-peaks per perturbation target, quantifying the differential regulation of highly variable peaks for each type of perturbation

These features were computed using the ArchR framework version 1.0.1 (Granja et al., 2021) with standard parameters unless otherwise stated. We provide each feature set as a dedicated h5ad file on scperturb.org, and our analysis roughly follows the pipeline proposed in Spear-ATAC (Pierce et al., 2021), as detailed below.

Note that these features were originally developed for scATAC-seq data on non-perturbed cells, with goals such as the identification of cell types, discovery of cell type-specific regulatory elements, or reconstruction of cellular differentiation trajectories (Buenrostro et al., 2013; Satpathy et al., 2019).

#### Pre-processing

Filtering out cells of low quality: To ensure a consistent and homogenous quality throughout the different data sets, we filtered out cells with fewer than 1000 and more than 100,000 mapped fragments. We further required a minimum transcription start site enrichment score of 4 to ensure a sufficient signal to noise ratio. See ArchR’s ‘createArrowFile’ function for details.

For the Spear-ATAC data set we ran ArchR’s getValidBarcodes function on processed 10x Cell Ranger files to subset the data set to valid barcodes. For the other datasets these files were unavailable, and we relied on the original authors’ pre-processing of barcodes.

Assigning sgRNAs to barcodes: For the Spear-ATAC and CRISPR_sciATAC datasets we had access to cell barcode-sgRNA count matrices (see original publications for details). We assigned the sgRNA with the highest counts to a cell barcode if the sgRNA count exceeded 20 and if that sgRNA combined at least 80% of all sgRNA counts. Cells that could not be assigned a sgRNA were left in the data set. For the ASAP-seq dataset a barcode-sgRNA matrix was not available. Instead, we relied on a sgRNA assignment downloaded from the study’s GitHub repository (Lareau, 2021).

#### Feature computation

All features described in the overview above were computed with ArchR functions. For details inspect the “fragments2outputs.R” script in our code repository (see Data Availability).

### scRNA-seq

#### Data acquisition

Datasets were downloaded from public databases following data availability directions in the source papers. When available from the authors, unnormalized pre-processed cell-by-gene matrices were used. Supplemental information from the papers were used in data analysis when applicable.

#### Data processing

Analysis started from unfiltered, unnormalized cell-by-gene matrices as provided by source papers. For one dataset, preprocessed cell-by-gene matrices were unavailable; pre-processing was performed following the procedure outlined in the original paper, directly using supplied code (Gehring et al., 2020). For datasets with cell barcodes, barcode assignments for cells were taken from the original paper when available; when not available, barcode assignment was performed as described in the methods section of the relevant paper. If multiple guides were assigned to the same cell, the guides were listed in decreasing order of counts in the final data object. The code used for processing each individual dataset, including barcode assignment, is available in our code repository.

Datasets were imported into AnnData objects using Scanpy (versions 1.7.2–1.9.1) (Wolf et al., 2018). Metadata was taken from the original papers when available. For cell lines, information on sex, age, disease, and origin were taken from Cellosaurus (Bairoch, 2018). Metadata columns are described in (Supp Table 2). Items listed in **bold** are included for all datasets.

Datasets are saved as .h5ad files and as .mtx files with obs and var as separate .csv files. Code is supplied in our code repository for the import of .mtx files into Seurat.

#### Data analysis

Before calculating the E-distances (Fig 4), cells and genes were filtered using Scanpy (versions 1.7.2–1.9.1) (Wolf et al., 2018). All .h5ad objects published on the resource were saved using Scanpy 1.9.1. Cells were kept if they had a minimum of 1000 UMI counts, and genes with a minimum of 50 cells. 2000 highly variable genes were selected using `scanpy.pp.find_variable_genes` with flavor ‘seurat_v3’. We normalized the count matrix using `scanpy.pp.normalize_total` and log-transformed the data using `scanpy.pp.log1p`; We did not z-scale the data. Next, we computed PCA based on the highly variable genes. The E-distances were computed in that PCA space using 50 components and Euclidean distance. To avoid problems due to different numbers of cells per perturbation, we subsampled each dataset such that all perturbations had the same number of cells. We removed all perturbations with fewer than 50 cells and then subsampled to the number of cells in the smallest perturbation left after filtering. Large parts of our analysis were parallelized as workflows using snakemake (Mölder et al., 2021).

#### E-distance

The E-distance is a statistical distance between high-dimensional distributions and has been used to define a multivariate two-sample test, called the E-test (Rizzo and Székely, 2016). It is more commonly known as energy distance, stemming from the original interpretation using gravitational energy in physics. Formally, it contextualizes the notion that two distributions of points in a high-dimensional space are distinguishable if they are far apart compared to the width of both distributions (Fig 3A). More specifically,

Let *x*_1_,…, *x*_*N*_ ϵ ℝ^*d*^ and *y*_1_,…, *y*_*M*_ ϵ ℝ^*d*^ be samples from two distributions *X,Y* corresponding to two sets of *N* and *M* cells, respectively.

We define

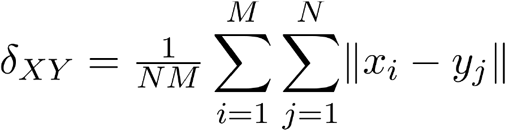

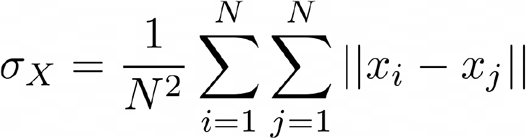

and defined σ_*y*_accordingly. We used the squared euclidean distance when calculating cell-wise distances. Intuitively, *δ*_*xy*_ is the mean distance between cells from the two distributions, while *σ*_*x*_ describes the mean distance between a cell *X* from *X*, to another cell from. The energy distance between *X* and *Y* is defined as:

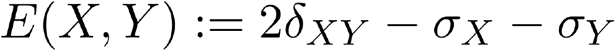

#### E-test calculation

The E-test was performed as a Monte Carlo permutation test using the E-distance as test statistic. For each dataset and each perturbation within that dataset, we took the cells and combined them with the unperturbed cells. Then, we shuffled the perturbation labels and computed the E-distance between the two resulting groups. We repeated this process 100 times. The number of times that this shuffled E-distance to unperturbed was larger than the unshuffled one divided by 100 yields a p-value, which we report for almost all datasets in our resource (Supp Table 3). We corrected for multiple testing using the Holm-Sidak method per dataset.

#### Cell type hierarchy

Cell type annotations celltype.l2 were used as provided by (Hao et al., 2021). Doublets were removed and data was subset to 91 cells per remaining cell type, which is the largest number such that all key cell types had at least that many cells. After subsetting, the data was processed as for other RNA datasets. Protein data was CLR normalized using Muon 0.1.2 and log-transformed prior to PCA (Bredikhin et al., 2022). The hierarchy was computed using `scipy.cluster.hierarchy.linkage` from scipy 1.8.0 with method “single”.

#### Subsampling analysis

At each subsampling point we computed detailed E-statistics (E-distances, delta, sigma, E-test results) from each perturbation to the corresponding unperturbed cells of that dataset using PCA with 50 components based on 2000 highly variable genes, except specified otherwise. We downsampled raw UMI counts using the function `scanpy.pp.downsample_counts` on raw counts, then preprocessed (normalized, log1p-transformed, etc.) the data as previously described. Cells were downsampled to the same number at each subsampling step across all perturbations to avoid comparability issues. If possible, we recalculated the PCA while keeping the highly variable genes originally obtained from the complete dataset. Figures 5C, 5D and Supplemental Figures 5A, 5B were computed as a running loss of E-test significance (p-value<0.05) of formerly – i.e. prior to any subsampling – significant perturbations while subsampling, then normalized across datasets through division by the total number of formally significant perturbations in that datasets.

#### Advice for single-cell perturbation analysis

Resource users should be aware that memory requirements quickly become a limiting factor, especially with the newer, larger datasets, such as ReplogleWeissman2022 with >2.5 million cells across more than 9000 perturbations (Replogle et al., 2022). For example, the E-distance presented here for calculating distances between perturbed sets of cells relies on principal component analysis (PCA), but computing PCA for all data in this dataset was not possible with 500GB of memory without modifications to accelerate computation. Going forward, computational methods will need to be modified as in (Dhapola et al., 2022) to reduce memory load, or datasets will need to be subsampled. Additionally, the .h5ad datasets shared in this resource can be programmatically accessed using .h5py, and perturbations of interest extracted without requiring full dataset access.

To our knowledge, there are not yet established best practices for analysis of single-cell perturbation data. DESeq2 is frequently used for differential expression testing, as it can be applied to pseudo-bulk profiles of each perturbation (Love et al., 2014). An optional next step would be enrichment analysis of the resulting genes. Averaging single-cell measurements over cells per perturbation simplifies analysis and reduces the effect of measurement noise significantly but comes at the cost of removing all system-intrinsic biologically relevant information in cell-to-cell variation. In many studies, these average profiles are then embedded using a dimensionality reduction method of choice and subsequently clustered to reveal groups of perturbations with potentially similar targets (Norman et al., 2019; Replogle et al., 2022).

## Funding / Acknowledgements

- National Resource for Network Biology (NRNB, P41GM103504)
- Supported by the Wellcome Leap ∆ Tissue Program
- Deutsche Forschungsgemeinschaft (DFG, RTG2424 CompCancer, Beyond the Exome)
- Einstein Stiftung Berlin (Einstein visiting fellow program)
- Computation was in part performed on the HPC for Research cluster of the Berlin Institute of Health.
- We appreciate informative conversations with Yuge Ji, helpful code suggestions from Garrett Wong, and computational support from Aaron Kollasch. We also appreciate preprint review comment from Arcadia Science’s preprint review initiative (Gregory P. Way, Natalie Davidson, Erik Serrano, Parker Hicks, Jenna Tomkinson, Dave Bunten).

## Supplement

**Supp Fig 1:**
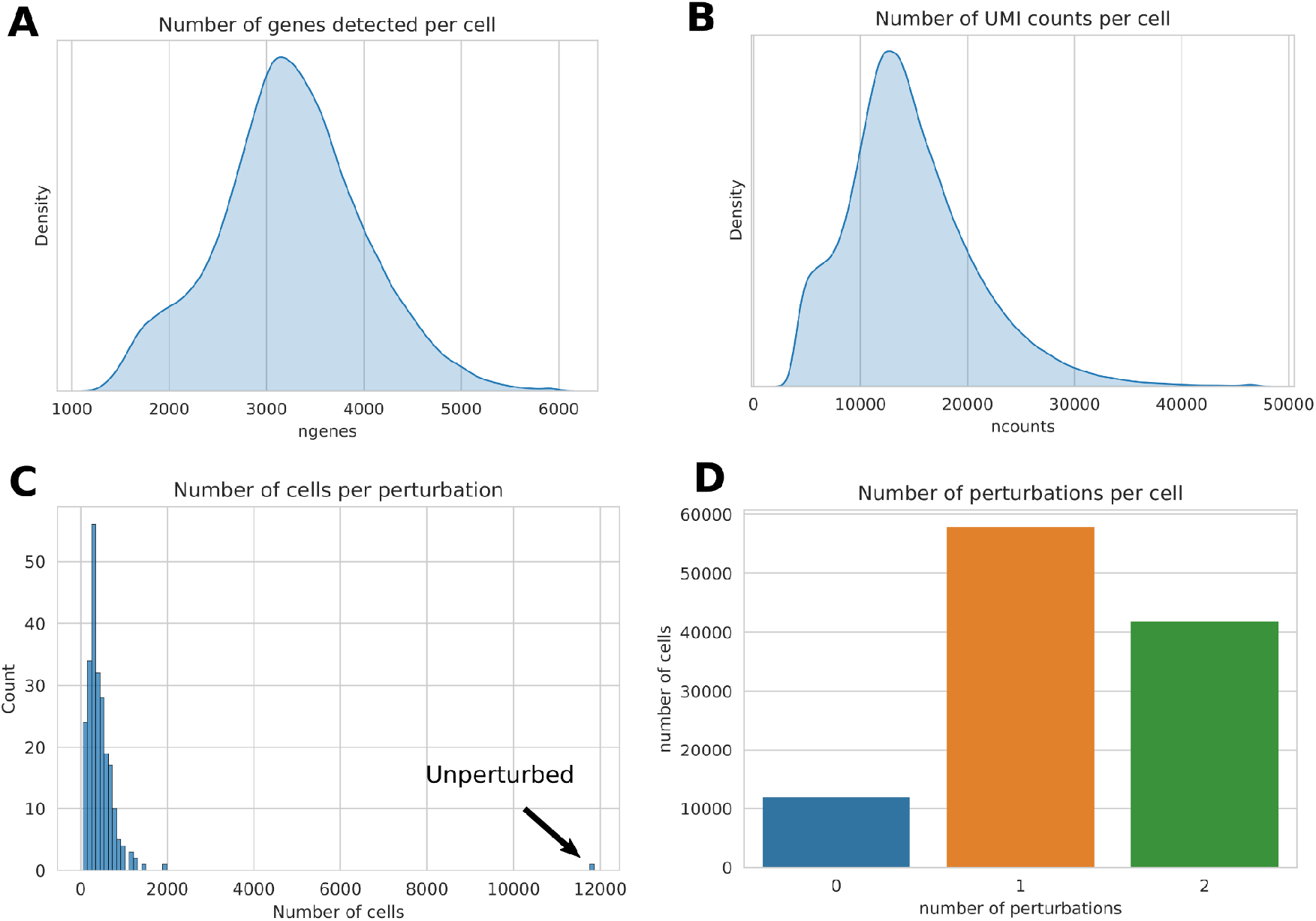
Exemplary information provided for each scPerturb dataset (here for (Norman et al., 2019)) (A) Number of genes that are detected with at least one count in a cell across all cells. (B) Total number of UMI counts per cell across all cells. Together with Supp Fig 1A this provides an overview over both sparsity and quality of the dataset. (C) Number of cells per perturbation. Depending on the application, perturbations with few cells can be filtered out before down-stream analysis. High imbalance in cell numbers per perturbation may also lead to biases in models. (D) Number of cells which received none, a single one, or two perturbations.

**Supp Fig 2A:**
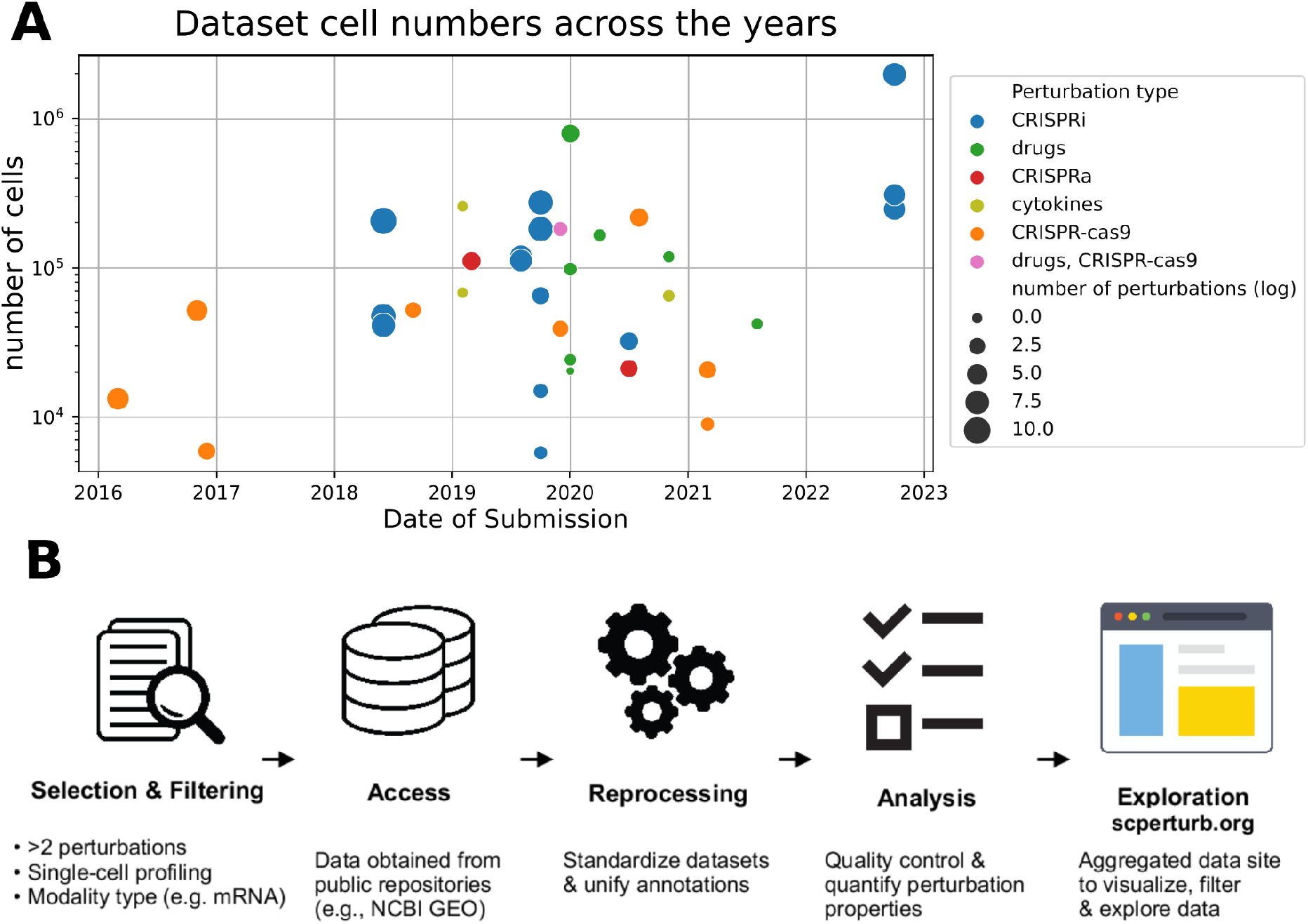
Number of cells per dataset by submission date. There is a rapid increase in published single-cell perturbation datasets around 2019. We speculate that the slight decrease of dataset numbers after 2021 suggested by the plot is due to the ongoing impact of reduced research in the earlier phases of the COVID-19 pandemic. **Supp Fig 2B: Harmonization and analysis workflow**. Perturbation datasets with single-cell molecular profiles with at least two perturbations and one control condition (e.g. unperturbed) of various modality types were identified in a literature search. Data was obtained from public repositories, and metadata (such as guide identity) from paper supplements. Datasets were reprocessed to standardize annotations and analyzed in parallel. All datasets are now available for download from scperturb.org, along with visualizations and summarizing information

**Supp Fig 3:**
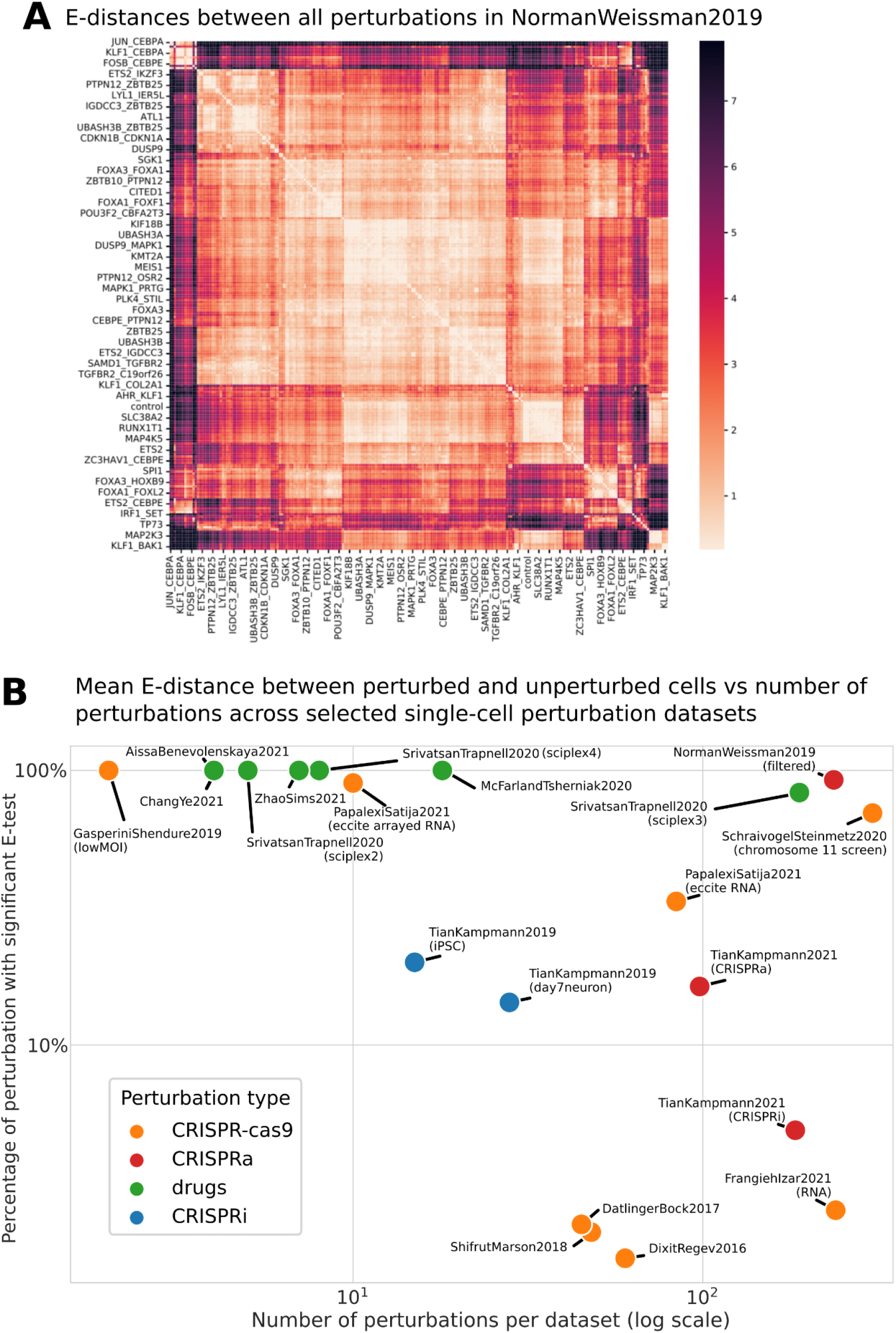
E-distance extended plots. (A) E-distances between all pairs of perturbations in the dataset NormanWeissman2019. The color scale is clipped at 5% highest and lowest percentiles. Clusters of similar perturbations are visible, e.g. a cluster of strongly acting perturbations targeting CEBPA at the top. (B) Percentage of perturbations with significant E-test to unperturbed cells in each dataset plotted against the total number of perturbations in the dataset (both in log scale).

**Supp Fig 5:**
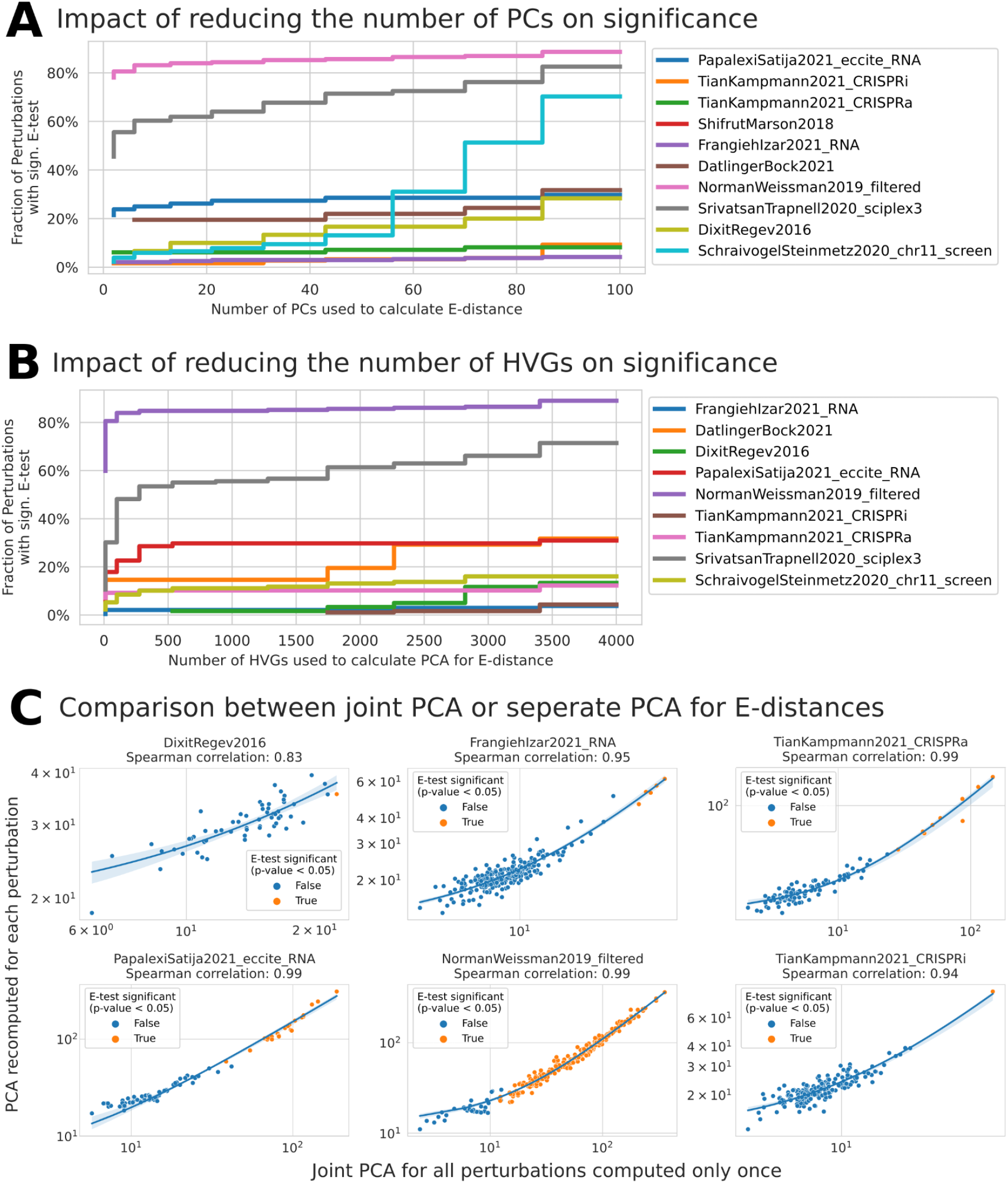
Tests on robustness of E-statistics to dataset properties and parameters. (A) Effect of using different numbers of principal components (PCs) from PCA on the number of perturbations with significant E-test w.r.t. unperturbed cells. The SchraivogelSteinmetz2020 dataset is TAP-seq, thus has much less genes measured than all other datasets. The faster decrease in significance observed in this dataset indicates stronger sensitivity on the number of PCs with fewer features available. (B) Effect of using different numbers of highly variable genes (HVGs) for the PCA calculation prior to E-testing. For most datasets, E-test results appear to stay comparable between 500 and 4000 HVGs. (C) E-distance computed in a single, joint PCA compared to E-distance computed in a separate PCA per perturbed-unperturbed combination across three exemplary datasets. Consistently high Pearson correlations indicate strong equivalence between both approaches across datasets.

